# Telomere-to-telomere gap-free genome assembly of a male donkey and the identification of novel SVs associated with functional genes

**DOI:** 10.1101/2025.09.12.675839

**Authors:** Tao Yang, Ran Yang, Yu Liu, Mo Feng, Yuanyuan Li, Hetong Zhang, Xinyu Wang, Ruyu Yao, Jiahui Wu, Wenhui Xing, Shiyu Qian, Chunjiang Zhao

## Abstract

Previous assemblies of the donkey genome remain with gaps and structural errors, and a complete donkey genome will greatly facilitate genetic research related to donkeys. In the present study, a 2.78-Gb telomere-to-telomere gap-free donkey genome (CAU_T2T_donkey) was assembled, including a 29.78-Mb Y chromosome, aided by ONT and trio-binning approach. CAU_T2T_donkey corrected the structural errors of previous assemblies and added a total of 153.8-Mb previously unresolved regions and 354 genes to the reference genome EquAss-T2T_v2. We identified a 1.9-Mb PAR on CAU_T2T_donkey-chromosome Y, and added 17.1Mb regions and 75 new genes to the chromosome Y of the previous reference genome ASM1607732v2. Multi-copy genes, such as *TSPY, L1RE, ETY, HSFY*, and *ETSTY* were also identified in CAU_T2T_donkey-chromosome Y. Totally 6 types of repetitive sequences in centromeric regions were identified, and the features of the centromeric regions were revealed, and satellite-free centromeres were identified. We aligned HiFi long-read sequences of donkeys from six breeds against CAU_T2T_donkey and identified SVs in previously unresolved regions, and some of the novel SVs were located in functional genes, such as *AOX1* (Chr4:DEL61), *ASIC2* (Chr13:INS954), and *Twist2* (Chr19:DEL98).

## Introduction

Donkeys, serving as a labor animal, have played as import role in agriculture and transportation. After its domestication about 6,000 years ago^1,2^, different genetic structures and breed-specific traits of domestic donkey populations were developed under nature and artificial selections. A high-quality donkey reference genome is essential for understanding the detailed chromosomal-level genetic information of this important livestock and revealing in-depth differences in genetic architecture across donkey breeds. Four assemblies of the donkey genome were released since 2015^3-5^, in which Wang et al. firstly assembled a chromosome-level genome (ASM1607732v2) in 2020, and a T2T assembly, EquAss-T2T, was released in 2024 and closed most gaps and corrected some assembly errors in ASM1607732v2. The previous donkey reference genomes provided powerful tools for genetic studies of donkeys. However, there are still some gaps in EquAss-T2T, mainly located in centromeric and telomeric regions, and some assembly problems remain unresolved. Moreover, the Y chromosome is absent from EquAss-T2T as it was derived from a female equid, while ASM1607732v2 only comprises a 12 Mb partial Y chromosome. All of the above limitations of previous assemblies emphasize the need for a complete donkey genome containing the Y chromosome.

The third-generation long-read sequencing technologies, mainly including PacBio HiFi and Oxford Nanopore Technologies (ONT), have brought new breakthroughs in genomic research^6,7^. With the aid of long-read sequencing technology, especially ONT, gap-free genome assemblies have already achieved for some plants^8-10^, human^11^, pig^12^, goat^13^, sheep^14^, and chicken^15^. The above achievement of complete assemblies has proven the effectiveness of long-read sequencing technologies in overcoming challenges that short-read technologies encounter, solving assembly issues related to centromeres, telomeres, and other complex regions of the genome, and eventually significantly enhancing genome completeness and accuracy^13,14,16^.

Donkey genome features for highly complicated centromeric regions, more acrocentric chromosomes, which impede the complete genome assembly in previous studies. In this study, we integrated trio-binning strategy with PacBio HiFi and ONT sequencing technologies to achieve a Telomere-to-Telomere (T2T) gap-free genome assembly of donkey (CAU_T2T_donkey). CAU_T2T_donkey encompasses all chromosomes, with a particular emphasis on centromeres, telomeres, and the Y chromosome. The new T2T assembly achieved in the present study resolved previously uncharacterized genomic regions, thus provides new insights into the structure and function of these regions. CAU_T2T_donkey offers a valuable resource for population genomics, variant detection, and genetic structure analysis, and significantly facilitates genetic research in donkeys.

## Results

### T2T gap-free genome assembly

A male individual (DZ240498) from Dezhou Donkey was selected for the genome assembly. We generated 256.0 (98.5×) Gb and 221.4 Gb (85.2×) of ultralong Oxford Nanopore Technology (ONT) reads and Pacific Biosciences (PacBio) HiFi reads, respectively, to assemble CAU_T2T_donkey (Supplementary Data 1). A total of 538.15 Gb of high-throughput chromatin capture (Hi-C) data (207.1×) and 71.60 Gb PromethION platform(Pore-C) data(27.5×) were used to scaffold the 330 contigs with an N50 of 79.78 Mb from the initial assembly, and anchor them to 32 pseudomolecules, including 30 autosomes and the two sex chromosomes (Figure1.a). The telomeric regions of the 32 chromosomes ranged from 1.19 to 30.20 kb. We eventually constructed a complete genome assembly of CAU_T2T_donkey with a size of 2.78 Gb, which was assembled from 32 scaffolds with a N50=105.15Mb (Table 1). Additionally, we also sequenced the genomes of the parents of the male donkey (DZ240498) with the Illumina NGS platform, and obtained 413.64 Gb and 330.01 Gb of short read data for the two donkeys, respectively. Chromosome Y was assembled with a trio-binning strategy.

**Table 1.**
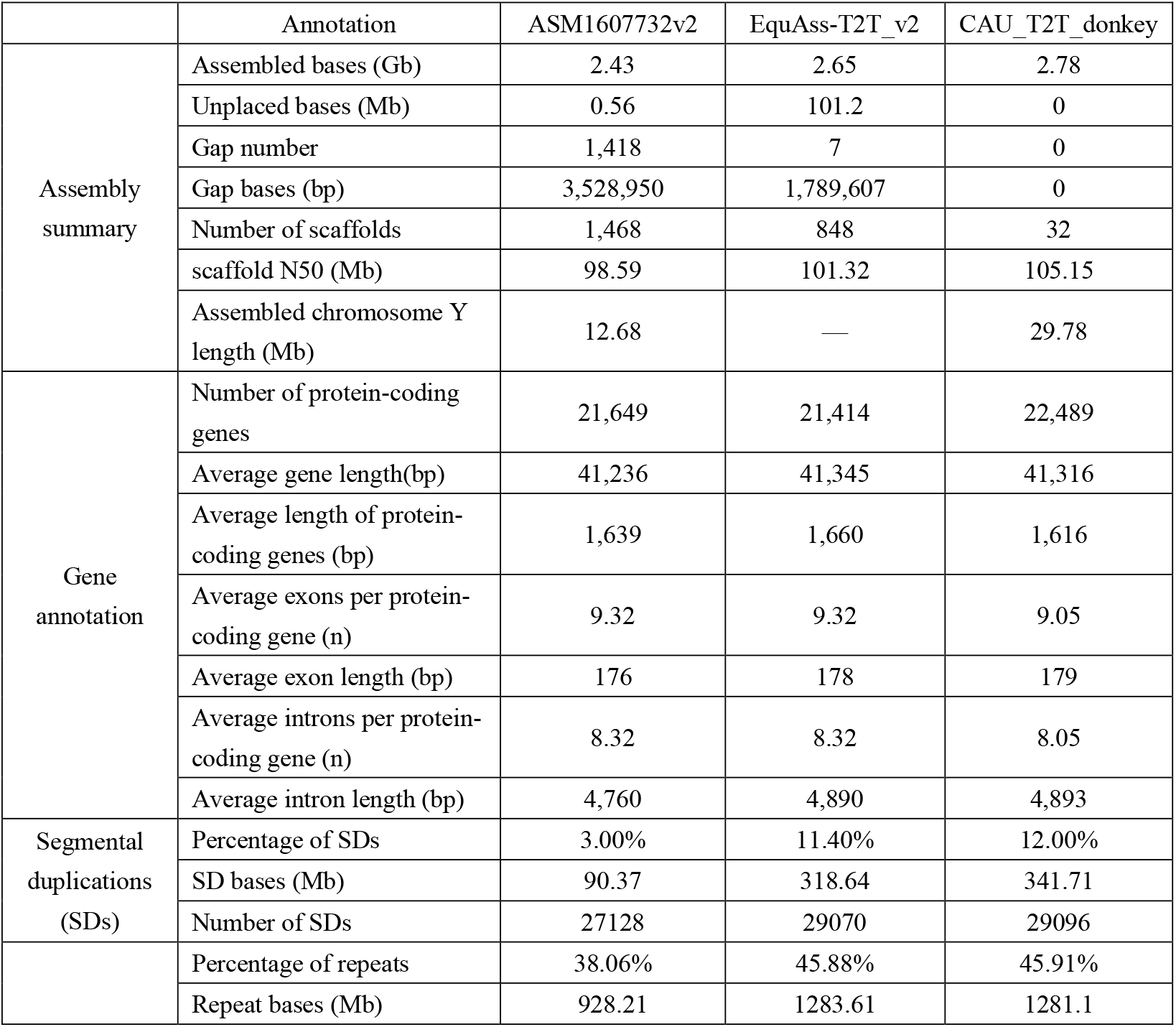
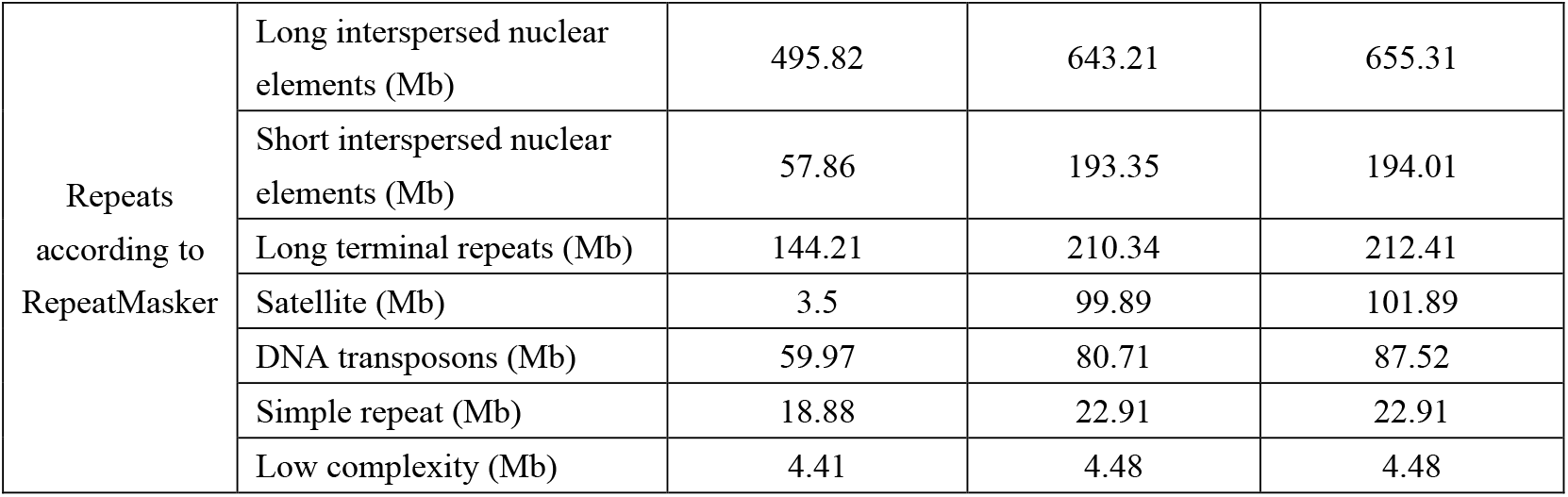
Comparison between CAU_T2T_donkey, EquAss-T2T_v2 and ASM1607732v2

### Validation and assembly improvement of CAU_T2T_donkey

Three strategies were applied to validate the quality of the CAU_T2T_donkey. The value of consensus base quality was 61.87(>99.99% base accuracy). The score of the phylogenetic coverage calculated with Benchmarking Universal Single-Copy Orthologs (BUSCO) based on laurasiatheria_odb10 was 98.80%. We aligned all of the long reads to CAU_T2T_donkey, and the even coverage of ONT and PacBio HiFi reads confirmed the reliability and the continuity of our assembly (Figure 1.b). Totally 64 telomeres were identified on 32 chromosomes (Supplementary figure 1), indicating a high-quality and complete telomere-to-telomere assembly of the CAU_T2T_donkey.

**Figure 1.**
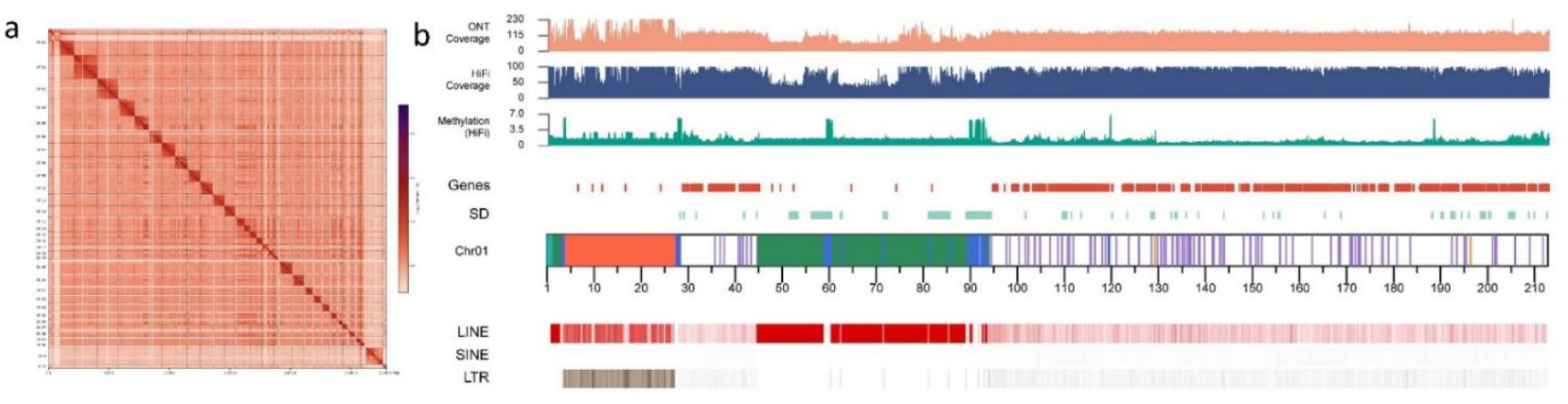
Assembly of CAU_T2T_donkey. a, pore-C heatmap for the chromosomes of the CAU_T2T_donkey assembly. All chromosomes were assessed with the pore-C heatmap. b, features of Chr01. The coverages of ultralong ONT and PacBio HiFi long reads are shown in 200-kb windows. The distributions of genes, SD, LINE, SINE, and LTR are shown in 200-kb windows using bars with different colors, respectively.

CAU_T2T_donkey showed a good collinearity with EquAss-T2T_v2 (GenBank accession GCA_041296235.2) (Figure 2.a). We filled the gaps left in EquAss-T2T_v2 and also corrected assembly errors in EquAss-T2T_v2, for example, the INV390 inversion found on Chr7 (Chr7: 79495463∼79698473). The alignment of the PacBio long reads validated that the region covering the INV390 was correctly assembled in CAU_T2T_donkey. Similarly, we also corrected another inversion error (INV 460, ChrX:134681763∼135102588) on ChrX of EquAss-T2T_v2 in our new assembly(Supplementary Figure 2).

**Figure 2.**
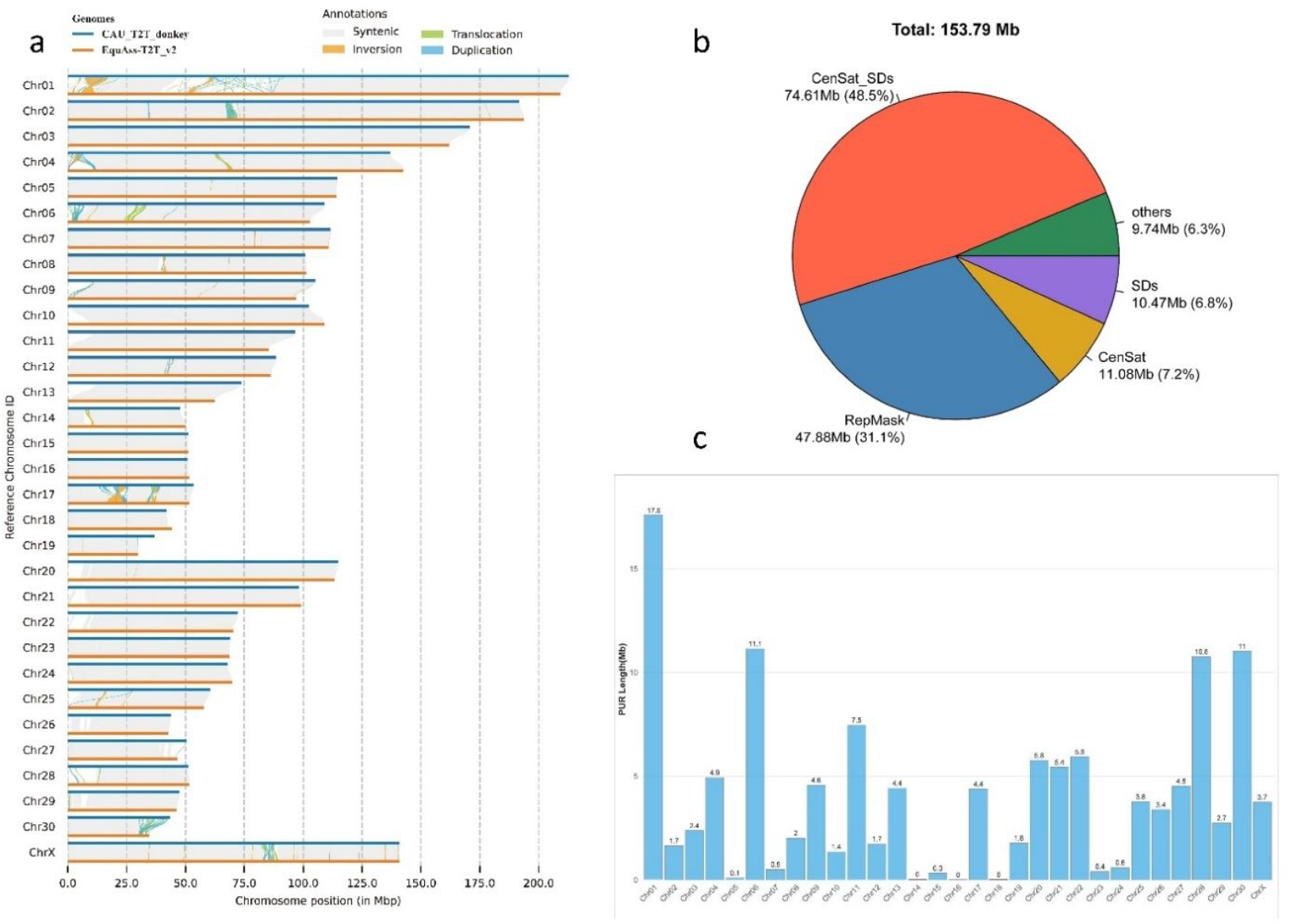
The assembly comparison between CAU_T2T_donkey and EquAss-T2T_v2. a, synteny between CAU_T2T_donkey and EquAss-T2T_v2. Blue line indicates CAU_T2T_donkey; yellow line, EquAss-T2T_v2. b, the proportions of various repetitive elements in PURs of EquAss-T2T_v2. CenSat, satellite sequences in the centromeric region; SDs, segmental duplications; RepMask, repeats by RepeatMasker; CenSat_SDs, bed regions with the overlapped SDs and centromeric satellite regions. c, the length of the PURs on each chromosome (excluding ChrY).

In EquAss-T2T_v2, there were 153.8 Mb unresolved regions (in the present study, the unresolved regions in previous assemblies are referred to as PURs), including unassembled and misassembled regions (Figure 2.b). Our new assembly shows that the PURs mainly contain centromeric segmental duplications (CenSat_SDs) (48.5%), RepMask (31.1%). Most PURs were distributed on Chr1, Chr6, Chr8, and Chr30, in which Chr1 harbored the greatest total length of PURs, spanning 17.6Mb (Figure 2.c). Some genes in PURs were annotated in CAU_T2T_donkey.

### Genome annotation

Repetitive sequences accounted for 45.91% (1,281.1Mb) of CAU_T2T_donkey, which was higher than that of EquAss-T2T_v2 (45.88% (1,263.83 Mb)) or ASM1607732v2 (38.06% (928.21Mb)) (Table 1). The majority of the repetitive sequences were retroelements, having a total length of 938.21Mb, accounting for 33.71% of genome. Abundant LINEs were identified in the genome with a total length of 655.31Mb and occupied 23.52% of the genome. We assembled satellite sequences with a total length of 101.89Mb, which accounted for 3.66% of CAU_T2T_donkey. The repetitive sequences of PURs occupied 11.19% of the whole repeats in the genome. Centromeric satellites were predominant in PURs and made up 48.5% of the sequences in PURs (Figure 2.b).

In total, 22,489 protein-coding genes were identified in CAU_T2T_donkey, of which 354 genes were located in PURs, including 40 newly annotated genes, and 222 genes were identified in centromeric regions. Additionally, we identified 29,096 SDs in CAU_T2T_donkey, and 15.15%(51.79 Mb) and 3.24% (11.08 Mb) of them were located in centromeric regions and PURs, respectively(Table 1,Figure 2.b).

### Centromeric regions and their repeat contents

We identified the positions of centromeres on the chromosomes based on the results of Chip-seq, heatmaps constructed with Hi-C or Pore-C data, and the position of enriched satellite sequences, as well as the binding regions of CENP-A reported previously (Cappelletti et al, 2025). We finally identified 14 metacentric or submetacentric chromosomes, and 18 telocentric or acrocentric chromosomes, in which ChrX is a metacentric chromosome and ChrY belongs to the acrocentric. The positions of the centromeres on the sex chromosomes are consistent with those reported previously^17,18^. We also found that the gaps left in EquAss-T2T_v2 were distributed in the centromeric regions of Chr14, Chr24, Chr26, and Chr28, which were filled in the assembly of CAU_T2T_donkey (Supplementary Data 2).

We identified totally 6 types of repetitive sequences in centromeric regions, which have repeat unit sizes of cirec1-114,cirec2-132, cirec3-221, cirec4-4171, and cirec5-4678, respectively, and LINE-1 was also observed in centromeric sequences. circ3-221 is a typical satellite DNA sequence found in equine species^19,20^, and it has the highest copy number in the genome in the satellites (Figure 3.b). Most chromosomes contain multiple repeat units. For example, Chromosome 13 includes cirec3-221, cirec1-114, cirec2-132, and cirec5-4678(Figure 3.a). Some chromosomes harbored a particular repeat unit type. For example, LINE-1 was highly enriched on Chr8, which was confirmed by fluorescence in situ hybridization(FISH) experiments (Figure 3.c). Similarly, four chromosomes (Chr4, Chr10, Chr12, and Chr30) carried 132 repeat units. Five chromosomes (Chr5, Chr7, Chr14, Chr16, Chr18, and Chr23) did not have any repeat units. Additionally, 45S rDNA is also a highly repetitive sequence. 6 chromosomes were harboring 45s rDNA, including Chr19, Chr20, Chr24, Chr25, Chr28, and Chr29, which were also confirmed with the results of FISH (Figure 3.c).

**Figure 3.**
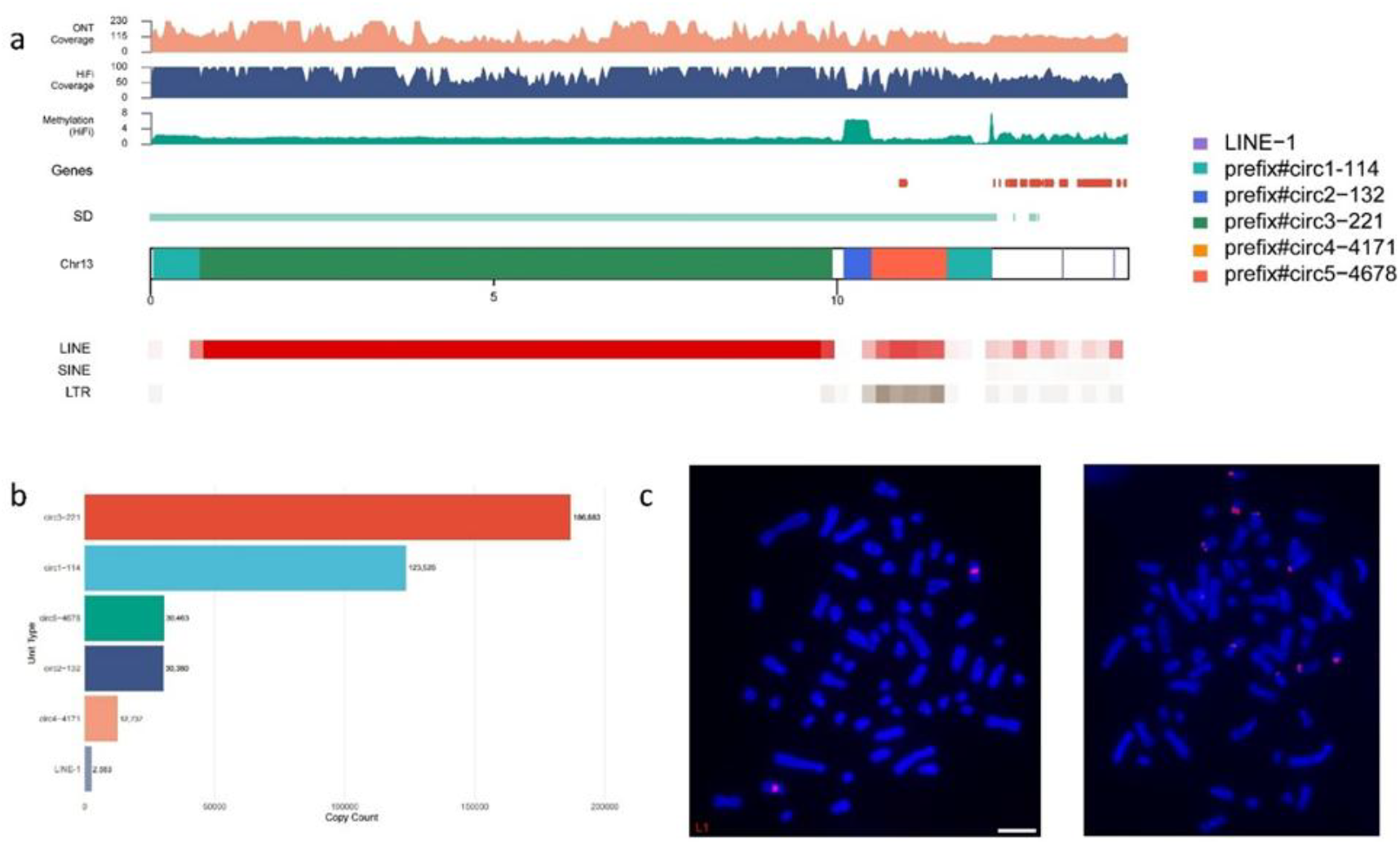
Distribution of the repetitive sequences on centromeric region and the 45S rDNA. a, the distribution of repetitive units on Chr 13. The various repeating units are indicated with different colors. The elements, such as genes, SDs, LINEs, SINEs, and LTRs are shown in 200-kb windows using bars with different colors, respectively. b, the copy numbers of each repetitive unit across the entire genome. c. FISH image of LINE-1 on Chr8 (left), with probes designed for both LINE-1 and Chr8 sequences; FISH image of 45S rDNA (right), with probes designed for conserved sequences of 45S rDNA (Supplementary Data 3).

### Y chromosome assembly and structure

As ChrY is abundant in repetitive sequences and shares PAR with ChrX, it is challenging to assemble the complete ChrY. In the present study, we assembled a 29.78-Mb complete donkey ChrY, which was 17.1 Mb longer than the previous assembly, ASM1607732v2-ChrY(Figure 4 a). There was a 1.9-Mb PAR on CAU_T2T_donkey-ChrY compared with the ChrX (Figure 4.b). A good collinearity was shown in the 3.45-Mb region between CAU_T2T_donkey-ChrY and ASM1607732v2-ChrY, and we identified some SVs at the end of ASM1607732v2-ChrY, including 2 INVs and 5 translocations (Figure 4.a).

**Figure 4.**
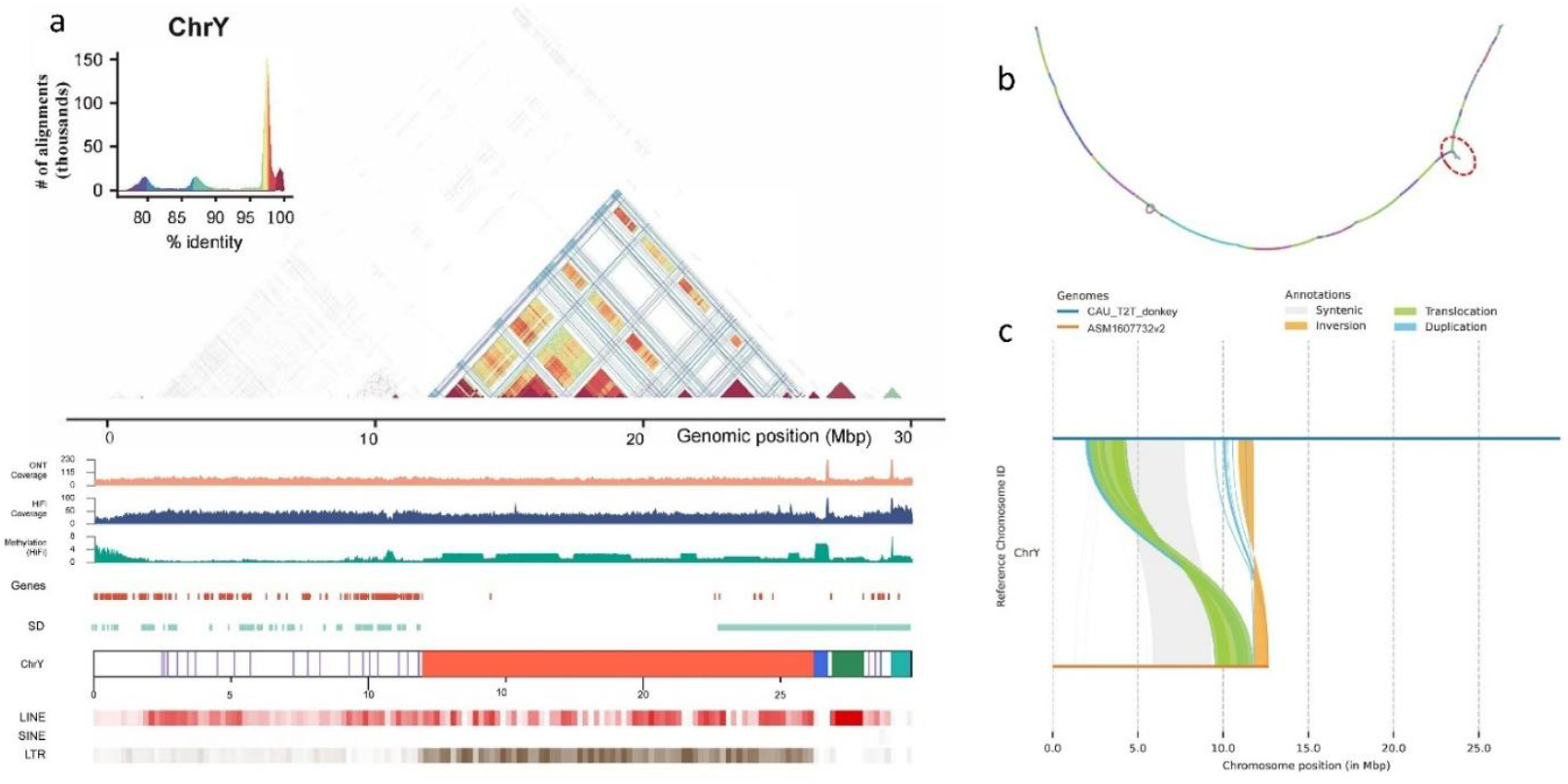
Assembly of chromosome Y. a, the features of CAU_T2T_donkey-ChrY, the elements such as genes, SDs, LINEs, SINEs, and LTRs are shown in 200-kb windows using bars with different colors, respectively. b, the tangle between the assemblies of ChrX and ChrY of CAU_T2T_donkey. c, the collinearity between CAU_T2T_donkey-ChrY and ASM1607732v2-ChrY.

The repetitive sequences on CAU_T2T_donkey-ChrY accounted for 77.72% of the chromosome, significantly higher than that of the whole genome (45.91%). There were abundant LTRs (10.57 Mb) and LINEs (10.32Mb) identified in the ChrY, and they accounted for 35.51% and 34.68% of the chromosome, respectively (Figure 4. d). The density of LTRs on the ChrY was much higher than that of the genome, in which LTRs only accounted for 7.61%. On the other side, we identified fewer DNA transposons (0.87%) and satellites (1.95%) on the ChrY than those of the whole genome, which were 3.14% and 1.95 %, respectively (Supplementary Data 4).

We identified 111 protein-coding genes, much more than the 65 annotated in ASM1607732v2-ChrY. 36 genes were annotated in both of the assemblies, for example, *SRY, Tmsb4x, Txlng*, and *XKR3. SRY* is well known as a gene for sex determination, while *Txlng* functions as binding protein encoded by Taxilinγ(a Y-linked pseudogene). Comparing with ASM1607732v2-ChrY, some multi-copy genes were identified in CAU_T2T_donkey-ChrY, such as *TSPY* (8 copies), *L1RE* (5 copies), *ETY* (4 copies), *HSFY* (3 copies), and *ETSTY* (3 copies). *TSPY* gene family and *ETSTY* are expressed tissue-specifically in testis, and the former involves in spermatogenesis, in which only *TSPY4* and *TSPY10* were identified in horse and donkey^21^. *HSPY* multi-copy gene family is unique to equines. We also annotated the gene *ZFY*, the coding gene for Zinc Finger protein.

### SV calling based on PacBio long reads

To assess the advantage of CAU_T2T_donkey as a reference for long-read alignment and calling of large-scale structural variations (SVs), we used the PacBio HiFi reads of six donkeys from six representative breeds with diverse origins. We identified a total of 45,618 SVs in the PacBio long reads of the six individuals (20,207∼22,073)using CAU_T2T_donkey as a reference (Figure 5.b, Supplementary Data 5). The predominant types of structural variants (SVs) were deletions (24,948) and insertions (20,154), while inversions, translocations, and duplications accounted for a small proportion. The sizes of deletions and insertions exhibited a distribution with a peak at 1–2 kb in length. A total of 4,112 core SVs were shared among all six individuals, while 18,961 SVs were unique to individual samples. These unique SVs are highly likely to be associated with breed-specific characteristics (Figure 5.a).

**Figure 5.**
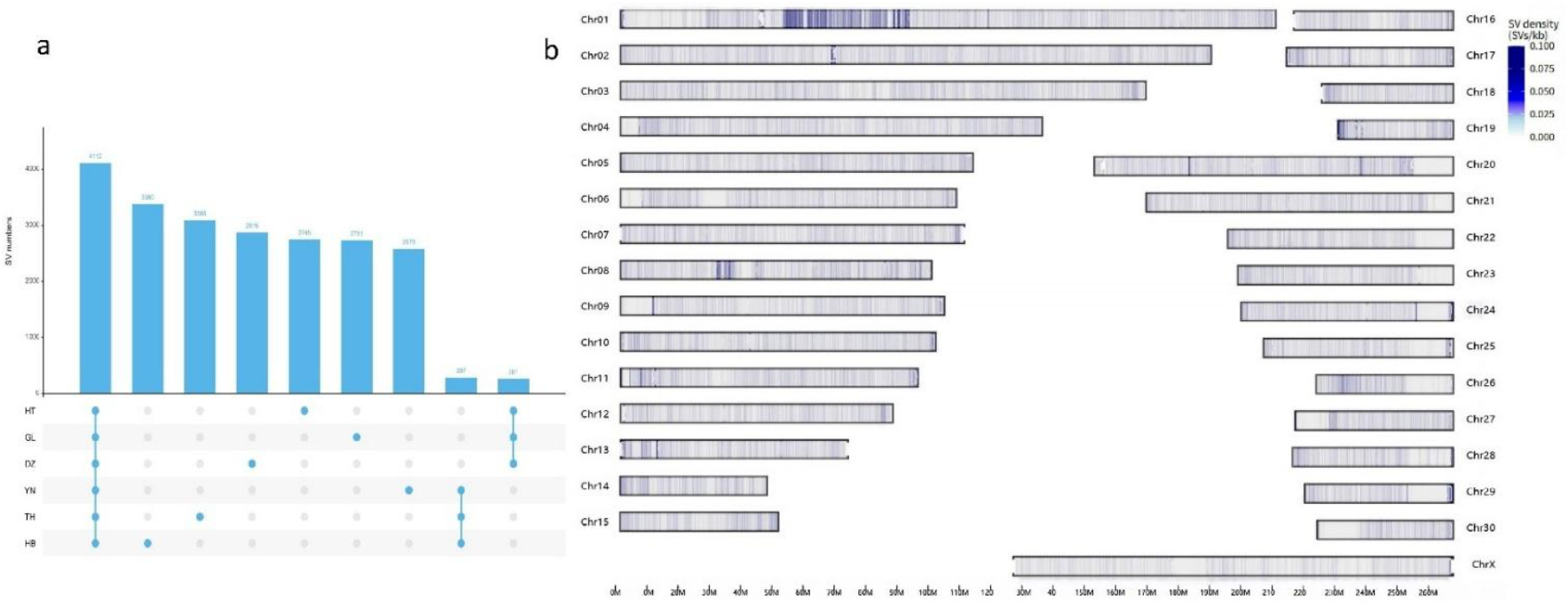
The structural variants screened with CAU_T2T_donkey. a, SVs uniquely identified in the six representative donkeys. HT indicates Hetian donkey; GL, Guangling donkey; DZ, Dezhou donkey; YN, Yunnan donkey; TI, Tibet donkey; TH, Taihang donkey. b, density of SVs on each chromosome called from PacBio reads of the six donkeys, in 10-kb windows.

The accurate assembly and completeness of the PURs in CAU_T2T_donkey enabled us to resolve complex genomic regions and enhance SV calling (Figure 5.b). Within the PUR region, 3,122 variants (6.84%) were identified, comprising 1,998 deletions and 1,124 insertions. We detected 14,482 SVs that fell into the gene bodies, with 121 SVs located within genes in the PUR region, including 47 SVs on ChrY (Supplementary Data 6). These novel variants were annotated to numerous functional genes, including *AOX1* (Chr4:DEL61), which is involved in muscle regeneration, lipid metabolism, and the regulation of fat-related traits in animals; *ASIC2* (Chr13:INS954) and Twist2 (Chr19:DEL98) were associated with traits of growth performance and body conformation. Interestingly, the insertion variant on *ASIC2* was identified exclusively in large body donkeys (Hetian donkey, Guangling donkey, and Dezhou donkey), while the deletion variant on Twist2 was detected only in small body donkeys (Yunnan donkey, Tibet donkey, and Taihang donkey). These findings suggest that these variants may influence donkey body size.

## Discussion

As the development of sequencing technologies, especially advent of the ONT sequencing, complete genome assemblies of several species have achieved^13,14,16^. The donkey is an important working animal that plays a vital role in agriculture and socioeconomic activities in many developing countries. Previous efforts to assemble high-quality donkey genomes have made considerable progress^3-5^. However, due to the limitations of sequencing technologies at the time, these assemblies still contained gaps and exhibited structural discontinuities, especially within highly repetitive regions such as centromeres and telomeres. In the present study, by integrating the latest sequencing technologies with a trio-binning strategy, we successfully generated a T2T gap-free donkey genome.

One of the remarkable improvements of the new assembly is that we assembled the complete donkey Y chromosome by overcoming the obstacle of complicated repetitive sequences and homology between sex chromosomes. The trio-binning strategy and ONT techniques enabled us to assemble a T2T gap-free ChrY. Compared with the previous ChrY assembly, we significantly improved the integrity and accuracy of the assembly of donkey ChrY. In the new assembly, we annotated novel genes missed in previous assemblies and more accurately determined copies of the important genes on ChrY, such as *TSPY, ETY, HSFY*, and *ETSTY*^3^, and the newly annotated genes and multi-copy genes have a crucial role in donkey reproductive capacity.

It is known that there are abundant telocentric and acrocentric chromosomes in donkeys, and it is also notable that the majority of the chromosomes harbor satellite-free centromeres (SFCs)^18^. In the present study, we successfully assembled the regions associated with telomeres and centromeres, which have long been considered major barriers to genome assembly. Previous studies showed that there are 16 donkey chromosomes carrying SFCs ^17,22^. It has been deduced that the SFCs were derived from the reposition of centromeres or Robertsonian translocation^18^. The SFC chromosomes are in the immature stage, and eventually satellites will accumulate in the centromeric regions and the centromeres will become stable^23^. Previous study showed that there were some repetitive sequences gradually accumulated in the centromeric regions of SFC chromosomes^3^, and our results also validated that. We found that the centromeric regions of 5 chromosomes in the donkey had almost no satellites or other repetitive sequences, while some other satellite-free centromeres, such as chromosome 8, were enriched with LINE-1 elements, which was consistent with the previous report that satellite-free centromeres of the Equus genus were rich in AT and LINE-1 sequences. These findings suggest that satellite-free centromeres are undergoing differential evolutionary trajectories. The availability of our complete telomere-centromere sequences provides valuable resources for elucidating the molecular mechanisms and evolutionary processes underlying centromere formation.

CAU_T2T_donkey enabled us to accurately and thoroughly screen SVs among breeds. The SVs within or neighboring genes were considered to be more likely to affect the genes’ function than SNVs. Novel SVs in PURs were identified with CAU_T2T_donkey, in which some SVs were harbored by functional genes related to growth and development, and notably, the SVs in the two genes *ASIC2* and *Twist2* were associated with body sizes. *ASIC2* is highly expressed in osteoblasts and can influence skeletal development and spinal^24-26^, and is also related to weaning weight and metabolic syndrome^27,28^. Previous study showed the *TWIST2* affected by 2q 37.3 interstitial deletion in human was associated with short stature^29^, and *TWIST2* was also related to the growth of skeletal muscle^30^. The identification of the novel SVs provides a good foundation for the study of gene functions.

In summary, we assembled the first T2T gap-free donkey genome in the present study. This high-quality assembly represents a significant improvement in both continuity and completeness compared with previous assemblies, and provides a critical genomic foundation for genetic research, conservation and utilization of donkey genetic resources. Furthermore, the fully resolved Y chromosome constitutes a key resource for studies on sex chromosome evolution, hybrid sterility, and male-specific traits in equids.

## Methods

### Ethical approval

All animal procedures were performed in accordance with the guidelines for animal experiments approved by the China Agricultural University Institutional Animal Care and Use Committee (AW13014202-1-1).

### Sample collection

For the genome assembly, peripheral blood samples of a 6-month-old male donkey (ID: DZ240498) and its parents from Dezhou donkey breed were collected at the Shandong Dong’e Black Donkey Breeding Center using BD Vacutainer EDTA tubes. Blood was then stored in a −80 °C freezer before the extraction of DNA or formaldehyde cross-linking for sequencing. High-molecular-weight genomic DNA was extracted based on the CTAB method, and purified with a Blood & Cell Culture DNA Kit (Cat# 13343, Qiagen, Beijing, China) following the manufacturer’s protocols. Sequencing for Pacific Biosciences (PacBio) reads, ultralong Oxford Nanopore Technology (ONT) reads, MGI (MGI Tech, Shenzhen, China) short reads, high-throughput chromatin capture (Hi-C) and PromethION platform data were all performed at the Benagen Ltd. (Wuhan, China).

In order to identify SV among different populations, six representative donkey breeds, including Guangling donkey, Dezhou donkey, Hetian donkey, Taihang donkey, Yunnan donkey, and Tibetan donkey, were selected and the blood of one animal from each breed was sampled in donkey preserving farms (Huabei grey donkey) or the preserving farms (other four donkeys) with the permission of their owners. The high-quality DNA was extracted from the five donkey samples as described above, respectively. And they were also sequenced for PacBio reads(15×) at the Benagen Ltd. (Wuhan, China).

### Fluorescence in situ hybridization (FISH)

Based on the sequences of centromere-specific satellites, fluorescently labeled probes were generated by PCR amplification using directly modified primers. Donkey bone marrow mesenchymal stem cells were cultured for preparing the slides with chromosomes at metaphase^31^. The metaphase chromosome slides were subjected to the following sequential treatments^32^: digestion with 1 ng/mL pepsin solution at 37 ℃ for 2 h, followed by fixation with 4% formaldehyde for 3 min. The hybridization mixture consisted of 50% formamide, 2× SSC, 10% dextran sulfate, and 100 ng of each probe. A total volume of 10 μL of the mixture was applied per slide. The probe in the mixture was denatured at 95 ℃ for 10 min and then incubated with the chromosome slides, which had been denatured separately in 70% formamide at 70 ℃ for 1 min. Hybridization was conducted at 37 ℃ for 18 h. Post-hybridization washes were performed as follows: three 5-min washes in 2× SSC, then a 5-min wash in 1× PBS. The slides were then counterstained with VectaShield antifade mounting medium containing DAPI. Finally, we obtained images with a DP80 camera integrated with an Olympus BX63 fluorescence microscope.

### Draft assembly and pseudo chromosome construction

Using hifiasm (0.25.0)^33^, we conducted a draft assembly with the long reads, including HiFi reads and ONT reads more than 50 K. Using the BLASTN tool (v.2.10.0), the primary assembly was searched against the nonredundant nucleotide database of NCBI to discard sequences of mitochondria and bacterial contaminants.

Based on the Pore-C sequencing data, we mapped all of the sequences to the 31 pseudo chromosomes (30 autosomes and X chromosomes) with the program cphasing (v0.7.1) using the following parameters: “(-n 31 –chimeric-correct --preset very-sensitive)”, and adjusted them with Jucerbox (v.2.18.0). Eventually, all scaffolds were anchored to the autosomes and ChrX.

### Initial assembly of ChrY

A trio-binning strategy was applied to assemble Chr Y. We conducted the haplotype assembly of the chromosome using verkk (v2.2) with the following parameters: “–telomere-motif CCCTAA –min-ont-length 50000 --porec --correct-min-read-length 3000 --correct-k-mer-size 1301 --hifi-coverage 90”. The initial assembly was aligned with ASM1607732v2 with minimap2, and only one gap was left in the initial T2T Chr Y assembly.

### Gap verification and filling

We conducted the gap verification by mapping the long reads of HiFi and ONT to pseudo chromosomes, and manually viewed the coverage of the reads with the Integrative Genomics Viewer (IGV) (v.2.13)^34^. To fill the gap, the ONT reads more than 50K were mapped to the bam files of the pseudo chromosomes or the initially assembled Chr Y. The reads without any hits or those that could be mapped with less than half of the aligned reads were screened, and contigs were assembled with the screened reads using NextDenovo (v.2.5.2)^35^. The contigs were then mapped to the initial assembly of the genome using minimap2—asm20, and the gaps were filled with the long reads or the contigs covering them.

### Telomere filling

The Telomeric regions were filled with three types of sequences. Type I was the HiFi reads harboring the telomere-specific repeats AACCCT/AGGGTT screened with BLASTN (v.2.10.0). All of the HiFi reads were mapped to the gap-less genome with Minimap2 (v.2.26)^36^and the reads that could not be aligned with the genome were classified as type II using SAMtools (v.1.18)^37^. The HiFi reads that could be aligned within 1-Mb region at the ends of chromosomes were defined as Type III. We used above three types of HiFi reads to perform the assembly of the telomeres with Hifiasm (0.25.0)^33^. After corrected, the contigs of the HiFi reads were scaffolded together with the sequences at the ends of each chromosome using RagTag(v.2.1.0)^33^. Sixty-four telomeric regions were merged at the two ends of 32 chromosomes.

### Genome polishing

We constructed 21-mer and 31-mer libraries with the short reads using yak (v0.1), and mapped the sequences of the libraries and HiFi reads to the gap-free genome using winnomap2(2.0.3). Then the genome assembly was polished with Nextpolish2 (v.0.2.0) with HiFi reads^39^. Finally, a gap-free complete genome assembly (CAU_T2T_donkey) of all chromosomes was constructed for donkey, with an average quality value of 61.87.

### Assessment and validation of genome assemblies

We validated the CAU_T2T_donkey genome assembly with coverage of reads, quality value, and collinearity with previous assemblies. Long reads were mapped to the new assembly using Minimap2 (v.2.26) and the Burrows-Wheeler Aligner(v.0.7.17), and depth coverage of the reads was determined in 200-kb windows using bamdst (https://github.com/shiquan/bamdst). The coverage was viewed with the R package karyoploteR (v.1.8.4)^40^. The collinearity between CAU_T2T_donkey and the previous assembly EquAss-T2T_v2 (GCF_041296235.1) was analyzed using Minimap2 (v.2.26) with the parameter “-cx asm10”. The synteny between them was plotted with paf2doplot (https://github.com/moold/paf2dotplot). We generated a 21-mer hash table with the short reads using Merqury (v.1.3.1)^41^ and determined the quality value and error k-mer frequency of CAU_T2T_donkey.

We determined BUSCOs with the laurasiatheria_odb10 database using BUSCO (v.4.0.5)^42^. All short reads were aligned to CAU_T2T_donkey assembly with the Burrows-Wheeler Aligner (v.0.7.17), and the mapping rate and base accuracy were calculated with SAMtools (v.1.18) and BCFtools (v.1.15.1).

### Annotation of repetitive sequences

We used de novo prediction and homolog-based methods to annotate the repetitive sequences. De novo prediction was conducted with EDTA2.2, and the repeats from de novo annotation were merged into a repetitive sequence database for homolog-based annotation using repeatMasker (v4.1.4) with default parameters^43^. We identified segmental duplications (SDs) with BISER80 (v1.4)^44^ and removed low-quality SDs according to the criteria described previously^13^.

### Protein-coding gene annotation

To annotate genes in CAU_T2T_donkey, we used following approaches, including homolog-based and transcriptome-based identification. To annotate with the homolog-based method, the homologous proteins of five species (donkey, horse, house mouse, pig, and human) were downloaded from NCBI, and they were aligned to CAU_T2T_donkey to identify the protein-coding genes using egapx(0.4.0) software^45^. For the transcriptome-based method, we downloaded RNA-seq data of 17 samples for 13 tissues of Dezhou Donkey and Iso-seq data for 14 tissues. We aligned the RNAseq data to our new assembly and conducted annotation using egapx(0.4.0). For the Iso-seq data, we acquired nonredundant transcripts using IsoSeq3 (v.3.8.2, https://github.com/PacificBiosciences/IsoSeq), and performed prediction of transcriptome using TransDecoder(v5.7.1). The annotation of EquAss-T2T_v2 was transformed with lifftoff. We integrated all of the gene prediction results using EVidenceModeler (v.1.1.1)^46^, and added alternative splicing transcripts and UTRs with PASA(v.2.5.2)^47^. At last, the gene names were added with Swissport(v0.11.2, https://github.com/swisspost).

### The determination of methylation using PacBio and short reads

The methylated sites of the genome were determined with HiFi and short reads. The HiFi reads were aligned to the CAU_T2T_donkey assembly using pbmm2 (v1.13.0) (https://github.com/PacificBiosciences/pbmm2) with default parameters. The prediction of methylation was conducted using pb-CpG-tools (v3.0.0) (https://github.com/Pacific-biosciences/pb-CpG-tools) with the parameter: aligned_bam_to_cpg_scores. The frequency of methylated cytosines was assessed in 20-kb windows using BEDTools (v.2.31.0). For the approach based on the short reads, quality control was performed using trim galore with the parameter: --clip_R1 5--three_prime_clip_R1 2--rrbs -o trimmed--basename. Then the short reads were aligned to the assembled genome, and methylated sites were extracted with MethylDackel (v1.11, https://github.com/dpryan79/MethylDackel) extract.

### Chip-seq and identification of centromeric sequences

The sequences of centromeres were extracted with the Chromatin immunoprecipitation (ChIP) assay with histone-specific antibodies. In the experiment, we treated fresh blood (10ml) of a Dezhou donkey with 1% formaldehyde for 10 minutes at room temperature and then quenched it by the addition of glycine (125 mmol/L final concentration). We performed immunoprecipitation with CENPA Rabbit pAb antibody (Cat # A15995, ABclonal, Wuhan, China). Immunoprecipitated DNA was used to construct sequencing libraries following the protocol provided by the I NEXTFLEX® ChIP-Seq Library Prep Kit for Illumina® Sequencing (NOVA-5143-02, Bioo Scientific) and sequenced on Illumina Novaseq 6000 with PE 150 method. Trimmomatic (version 0.36) was used to filter out low-quality reads^48^. Clean reads were mapped to the CAU_T2T_donkey. Genome by Bwa (version 0.7.15) ^49^. Samtools (version 1.3.1) was used to remove potential PCR duplicates[4].MACS2 software (version 2.1.2) was used to call peaks by default parameters (bandwidth,300 bp; model fold, 5, 50;p value, 0.001). If the summit of a peak is located closest to the TSS of one gene, the peak will be assigned to that gene^50^. HOMER (version 4.11) was used to predict motif occurrence within peaks^51^.

### Repeat identification in centromeres

The sequences of centromeric regions were identified with heatmaps constructed with Hi-C data and the ChIP-seq peaks. Construction of k-mer libraries was conducted using KMC(v3.1.1) with parameter”-fm -k151 -ci20 -cs100000”^52^. Centromeric repeats were retrieved with Satellite Repeat Finder based on the k-mer and the frequency of the k-mer in the region of centromeres. The repetitive sequences in the centromeric regions were aligned with CAU_T2T_donkey, and their abundance was estimated with BLASTN (v2.10.0).

### Structural variation identification

To obtain a high-quality collection of structural variants (SVs), we utilized four tools for the SV calling, namely, pbsv (v2.9.0, https://github.com/PacificBiosciences/pbsv), cuteSV (v2.0.2)^53^, svim(v2.0.0, https://github.com/eldariont/svim), and Sniffles (v2.0.7, https://github.com/fritzsedlazeck/Sniffles). We first aligned the HiFi reads to CAU_T2T_donkey to generate bam files using Minimap2 and SAMtools (v1.16). With the program pbsv, we investigated SV signatures based on bam files and called SVs based on PacBio reads for all six donkey samples using the “discover” and “call” modules of pbsv. For the programs cuteSV、 Sniffles, and svim, we performed SV calling with default parameters and added the --minsvlen 30. The SVs identified by the four programs were merged using the “merge” command of SURVIVOR^54^.

## Acknowledgements

This study was financially supported by the Key Research and Development Project of College and Enterprise, the Study on Comprehensive Supporting Technologies for Breeding of Dong-e Black Donkey (Grant No. 201605410411094), the Program for Changjiang Scholars and Innovative Research Team in University (Grant No. IRT1191), the Project on the Third National Survey of the Livestock and Poultry Genetic Resources (Grant No. 19221073), the project of the Beijing Key Laboratory for Genetic Improvement of Livestock and Poultry (Grant No. Z171100002217072), and the project of the National Germplasm Center of Domestic Animal Resources.

## Data Availability

The genome assemblies of CAU_T2T_donkey are uploaded to the NCBI database under bioProject accession PRJNA1316099 (https://www.ncbi.nlm.nih.gov/bioproject/PRJNA-1316099)

